# Vulnerability in the breadth evolution of an influenza broadly neutralizing antibody

**DOI:** 10.1101/2025.11.03.686356

**Authors:** Katrine E. Dailey, Yiquan Wang, Qi Wen Teo, Chaoyang Wang, Ruipeng Lei, Meixuan Tong, Lucia A. Rodriguez, Letianchu Wang, Nicholas C. Wu

**Author notes:** To whom correspondence may be addressed. (N.C.W.).

## Abstract

The highly conserved influenza hemagglutinin (HA) stem domain is a major target for broadly neutralizing antibodies (bnAbs). However, despite being discovered more than a decade ago, the IGHV1-69-encoded CR9114 remains the only HA stem bnAb that cross-reacts with both influenza A and B viruses. To investigate the constraints on the breadth evolution of CR9114, this study performs four deep mutational scanning experiments to compare the binding affinity landscapes of the germline and somatic CR9114 against H1 HA, H3 HA, and influenza B HA. Many mutations that minimally affect or even improve the H1 HA binding are detrimental for binding to H3 HA and BHA. We further reveal the prevalence of epistasis in IGHV1-69 HA stem bnAbs. Overall, our findings provide a mechanistic explanation for the scarcity of HA stem bnAbs with cross-reactivity against both influenza A and B viruses, and have important implications for developing broadly protective influenza vaccines.

## INTRODUCTION

Influenza viruses have four types (A-D), with types A and B being responsible for global public health concern. Hemagglutinin (HA), which consists of a hypervariable head domain atop a conserved stem domain^1^, is the major surface antigen of influenza A and B viruses. Influenza A HA is categorized into 19 subtypes (H1-H19) that fall into two groups. Group 1 includes H1, H2, H5, H6, H8, H9, H11, H12, H13, H16, H17, H18 and H19, whereas group 2 includes H3, H4, H7, H10, H14, and H15^2^. Similarly, influenza B HA (BHA) has two lineages, Yamagata and Victoria, although the Yamagata lineage has not been detected since the COVID-19 pandemic^3^. Each year, seasonal influenza epidemic, which is caused by H1 and H3 subtypes of influenza A virus as well as influenza B virus, leads to widespread illness, absenteeism, and loss of life^4^. Additionally, influenza A virus has caused five pandemics in the past 150 years with a high mortality burden, and remains at high risk for causing a future pandemic^5–8^.

Seasonal influenza vaccines, which are designed to elicit neutralizing HA antibodies, are currently the most effective preventive measures against influenza viruses. However, despite annual updates, the rapid antigenic drift of HA in seasonal influenza viruses sometimes leads to antigenic mismatch between the vaccine and circulating strains, resulting in low vaccine effectiveness in some years^9^. Furthermore, seasonal influenza vaccines do not confer protection against zoonotic subtypes that are antigenically distinct from human strains^10^. The limited heterosubtypic protection of seasonal influenza vaccines is largely attributable to the antibody responses being mostly directed against the hypervariable head domain, which is immunodominant over the conserved stem domain^11^. As a result, a major goal of the influenza research field is to develop more broadly protective vaccines.

HA stem has been the major target for the development of broadly protective influenza vaccines, owing to the discovery of broadly neutralizing antibodies (bnAbs) to the HA stem almost two decades ago^12–14^. Among hundreds of HA stem bnAbs that have been identified and characterized, CR9114 has the largest cross-reactivity breadth^15^. CR9114 not only binds to both group 1 and 2 HAs, but also BHA^15^. By contrast, none of the other known HA stem bnAbs have demonstrated the ability to cross-react with both influenza A and B HAs. The rarity of HA stem bnAbs exhibiting influenza A and B cross-reactivity may be attributed to the as-yet-unknown constraints of the evolutionary pathways required for their breadth evolution. By focusing on 16 somatic hypermutations in CR9114, a previous study has shown that the inferred germline of CR9114 (hereinafter referred to as germline CR9114) is specific to H1 HA and subsequently evolved breadth by acquiring affinity to H3 HA via somatic hypermutations, then to BHA^16^. However, it is unclear how the ability of CR9114 to evolve such exceptional breadth would be impacted by an alternative evolutionary pathway involving other potential somatic hypermutations.

In this study, we performed a series of Tite-Seq^17^ experiments to quantify the effects of almost all possible single amino acid mutations in the heavy chain variable domain (V_H_) of the germline and somatic CR9114 against different HAs. We found that the mutational tolerance of somatic CR9114 was the highest against H1 HA and lowest against BHA. Consequently, a number of mutations neutral or beneficial to germline or somatic CR9114 against H1 HA could abolish binding to H3 HA and BHA. These findings suggested that the evolution of broad reactivity by CR9114 is highly constrained and easily derailed due to epistasis and pleiotropy. Throughout this study, antibody residues are numbered using Kabat nomenclature.

## RESULTS

### Deep mutational scanning of the germline and somatic CR9114

To probe the evolutionary landscape of CR9114, we aimed to determine the binding affinity of all possible single amino acid mutations of the germline and somatic CR9114 against different HAs (**Figure 1**). Briefly, site-saturation mutagenesis was performed on the heavy chain variable domain of the germline and somatic CR9114. We did not include any mutations in the light chain, since it is not involved in binding^15^. Next-generation sequencing indicated that the frequencies of the mutations were fairly even (**Figure S1**). The mutant libraries were then displayed on the yeast surface and selected for binding against different HAs. Since germline CR9114 does not cross-react with group 2 HAs and BHA^16^, the mutant library of the germline CR9114 was selected against a recombinant HA stem construct that was designed based on H1N1 A/Brisbane/59/2007 HA (hereinafter referred to as H1 stem)^18^. On the other hand, the mutant library of somatic CR9114 was selected against H1 stem, H3N2 A/Singapore/INFIMH-16-0019/2016 (H3/Sing16) HA, and influenza B/Lee/1940 (B/Lee40) HA. The apparent binding affinity (K_D,app_) of each mutation was then quantified by Tite-Seq^17^, which combined fluorescence-activated cell sorting (FACS) and next-generation sequencing (**Figure 2 and Table S1**). For each mutation, we used - Δlog_10_ K_D,app_, to quantify its effect on binding affinity. The -Δlog_10_ K_D,app_ value represented the difference between the Δlog_10_ K_D,app_ of a given mutation and that of the corresponding wild type (i.e. germline or somatic CR9114). A lower -Δlog_10_ K_D,app_ value indicated a more detrimental effect on binding affinity. Pearson correlation coefficients of 0.94 to 0.99 were observed between the - Δlog_10_ K_D,app_ values from two replicates of Tite-Seq experiments, demonstrating the high data quality (**Figure S2A**). As a control, we also measured the expression level of each mutation by FACS and next-generation sequencing. Our result showed that most mutations have high expression level, which did not correlate with binding affinity (**Figure S2B**). Moreover, the binding affinity and expression level of nonsense mutations were distinct from those of silent mutations (**Figure S2B**), further validating our data.

**Figure 1.**
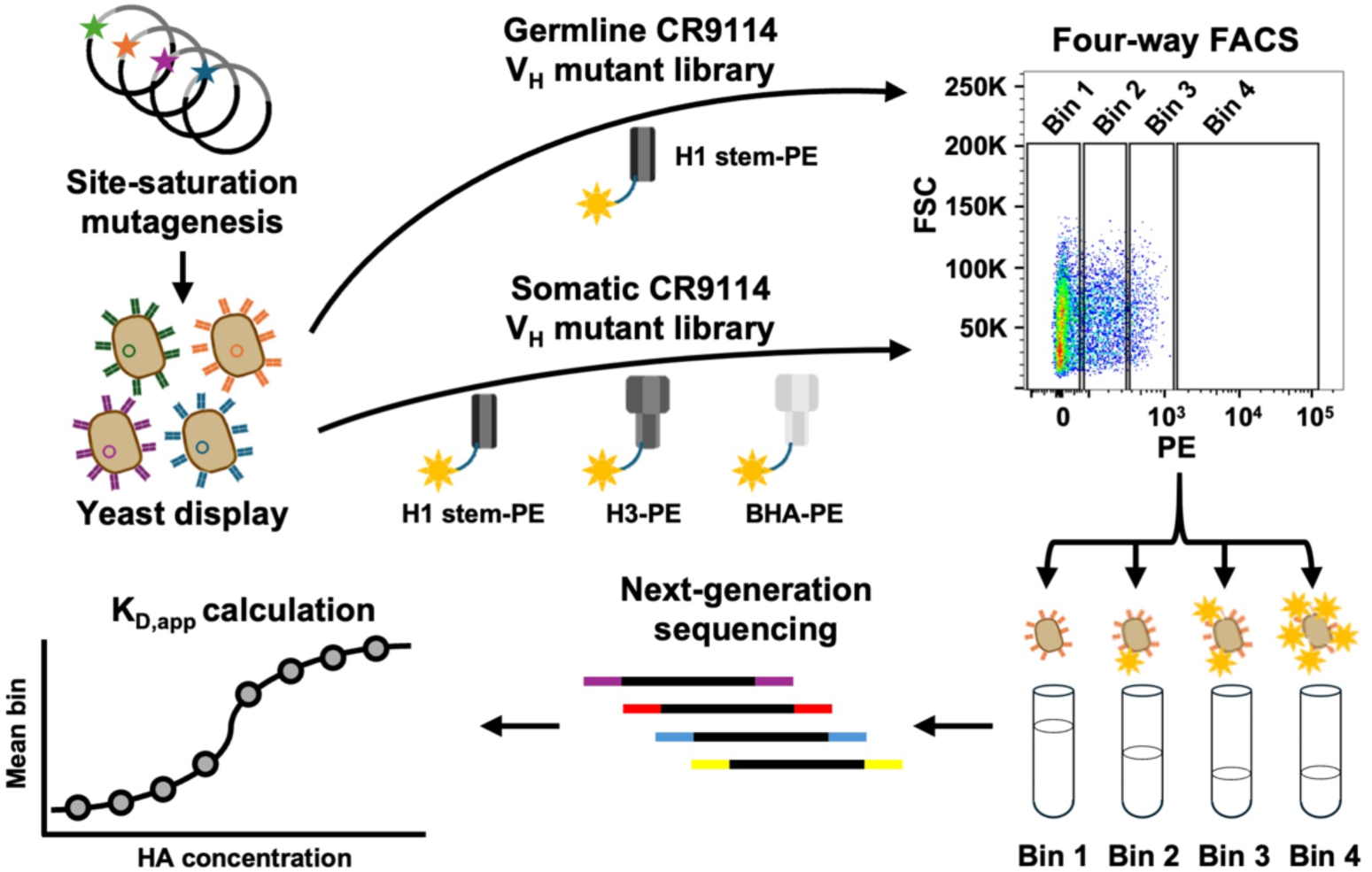
Workflow summary. A schematic representing our Tite-Seq experiments is shown.

**Figure 2.**
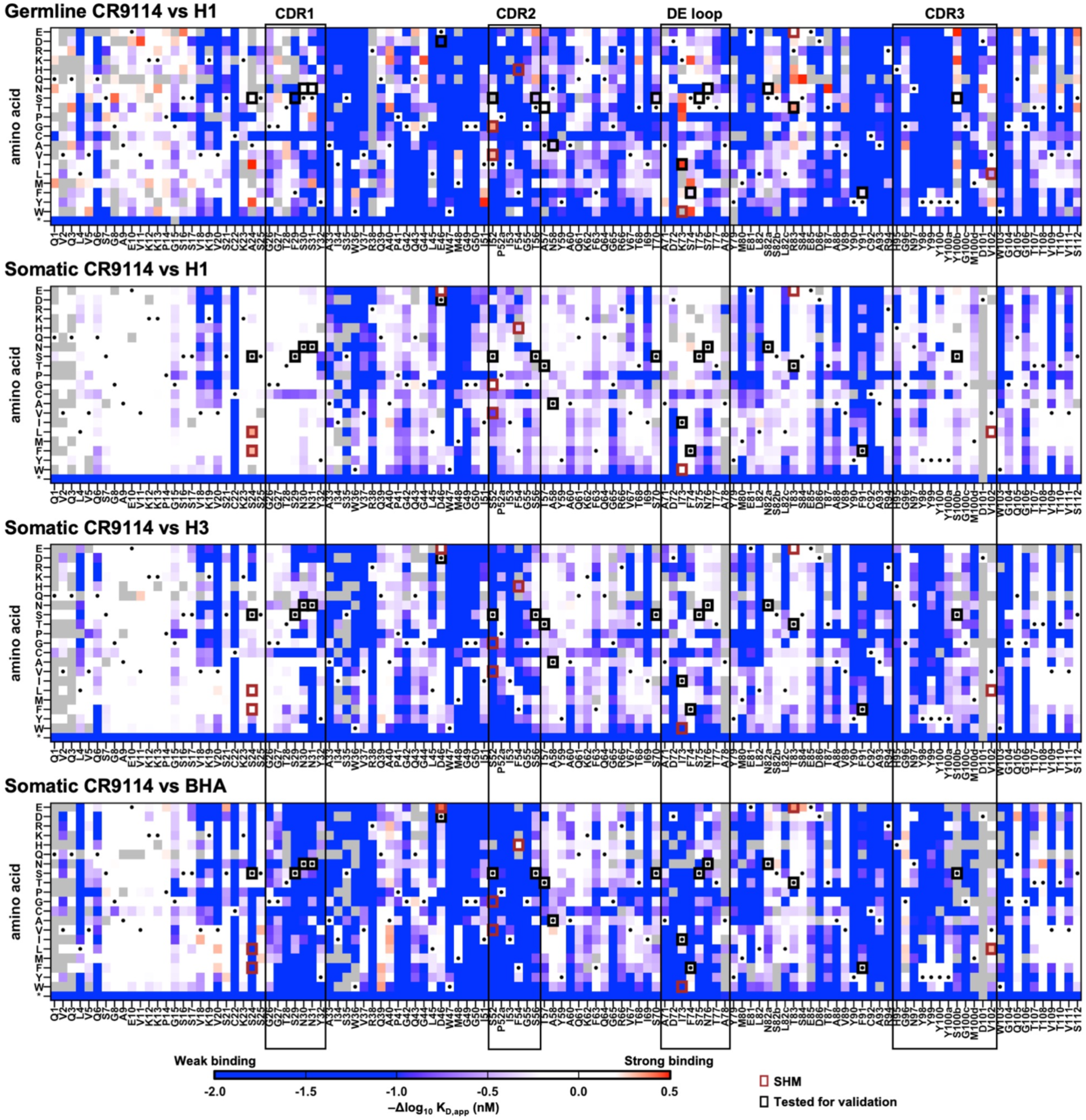
Binding affinity landscapes of germline and somatic CR9114 heavy chain. **(A-D)** Heatmaps showing the -Δlog_10_ K_D,app_ values of **(A)** single mutations of germline CR9114 against H1 stem, **(B)** single mutations of somatic CR9114 against H1 stem, **(C)** single mutations of somatic CR9114 against H3/Sing16 HA, and **(D)** single mutations of somatic CR9114 against B/Lee40 HA. Grey tiles indicate mutations that were discarded for our analysis due to low occurrence frequency (**see Methods**). Black rectangles spanning all four heatmaps indicate the locations of the CDRs and DE loop. Black dots indicate the corresponding wild-type sequence (i.e. germline or somatic CR9114). Black squares indicate the somatic hypermutations in CR9114. Brown squares indicate the mutations that were included in our validation experiment. “H1”, “H3”, and “BHA” indicate H1 stem, H3/Sing16 HA, and B/Lee40 HA, respectively.

### Mutational tolerance of CR9114 correlates with binding affinity

A previous study demonstrated that the binding affinity of somatic CR9114 to H1 HA was stronger than its affinity to H3 HA, which in turn was stronger than its affinity to BHA^16^. Consistently, our biolayer interferometry experiments showed that somatic CR9114 fragment antigen-binding (Fab) bound to H1 stem, H3/Sing16 HA, and B/Lee40 HA with K_D_ values of <0.1 nM, 36 nM, and 110 nM, respectively (**Figure S3 and Figure S4**). This order of binding affinity of somatic CR9114 against different HAs appeared to correlate with the mutational tolerance, which decreased from H1 stem (median -Δlog_10_ K_D,app_ = -0.22) to H3/Sing16 HA (median -Δlog_10_ K_D,app_ = -0.40) to B/Lee40 HA (median -Δlog_10_ K_D,app_ = -0.82, **Figure 2 and Figure S5**). Similarly, germline CR9114 had a weaker binding affinity (K_D_ = 185 nM) and lower mutational tolerance (median -Δlog_10_ K_D,app_ = -0.85) than somatic CR9114 against H1 stem (**Figure 2, Figure S3, and Figure S5**). It is well established that proteins with higher thermostability have higher evolvability ^19^. Nevertheless, despite the tendency of affinity maturation to decrease antibody thermostability^20^, the thermostabilities of germline CR9114 (T_m_ = 77.5°C) and somatic CR9114 (T_m_ = 78.5°C) were similar (**Figure S6**). Consequently, binding affinity seemed to play a more important role than thermostability in determining the mutational tolerance of CR9114.

### Prevalence of epistasis and pleiotropy in the breadth evolution of CR9114

The binding affinity landscapes of the germline and somatic CR9114 against different HAs allowed us to analyze the involvement of epistasis and pleiotropy in the breadth evolution of CR9114. Here, epistasis refers to how a given mutation has different effects on binding affinity between germline and somatic CR9114, whereas pleiotropy describes how a given mutation affects the binding affinity against different HAs. We first observed that mutations with minimal effects on the binding affinity of germline CR9114 to the H1 stem rarely affected the binding affinity of somatic CR9114 to H1 stem (**Figure 3A and Figure S7A**), but were frequently detrimental to the binding affinity of somatic CR9114 to H3/Sing16 HA (**Figure 3B and Figure S7B**), and even more often deleterious to that of B/Lee40 HA (**Figure 3C and Figure S7C**). These increasingly detrimental mutations throughout the screens mostly resided within the complementarity-determining regions (CDRs), while the effects of mutations in the framework regions on binding affinity correlated well between germline and somatic CR9114 (**Figure 3A-C and Figure S7A-C**). Our findings indicated that affinity maturation of germline CR9114 against H1 HA could impose a genetic barrier for evolving cross-reactivity to H3 HA and BHA due to epistasis and pleiotropy.

**Figure 3.**
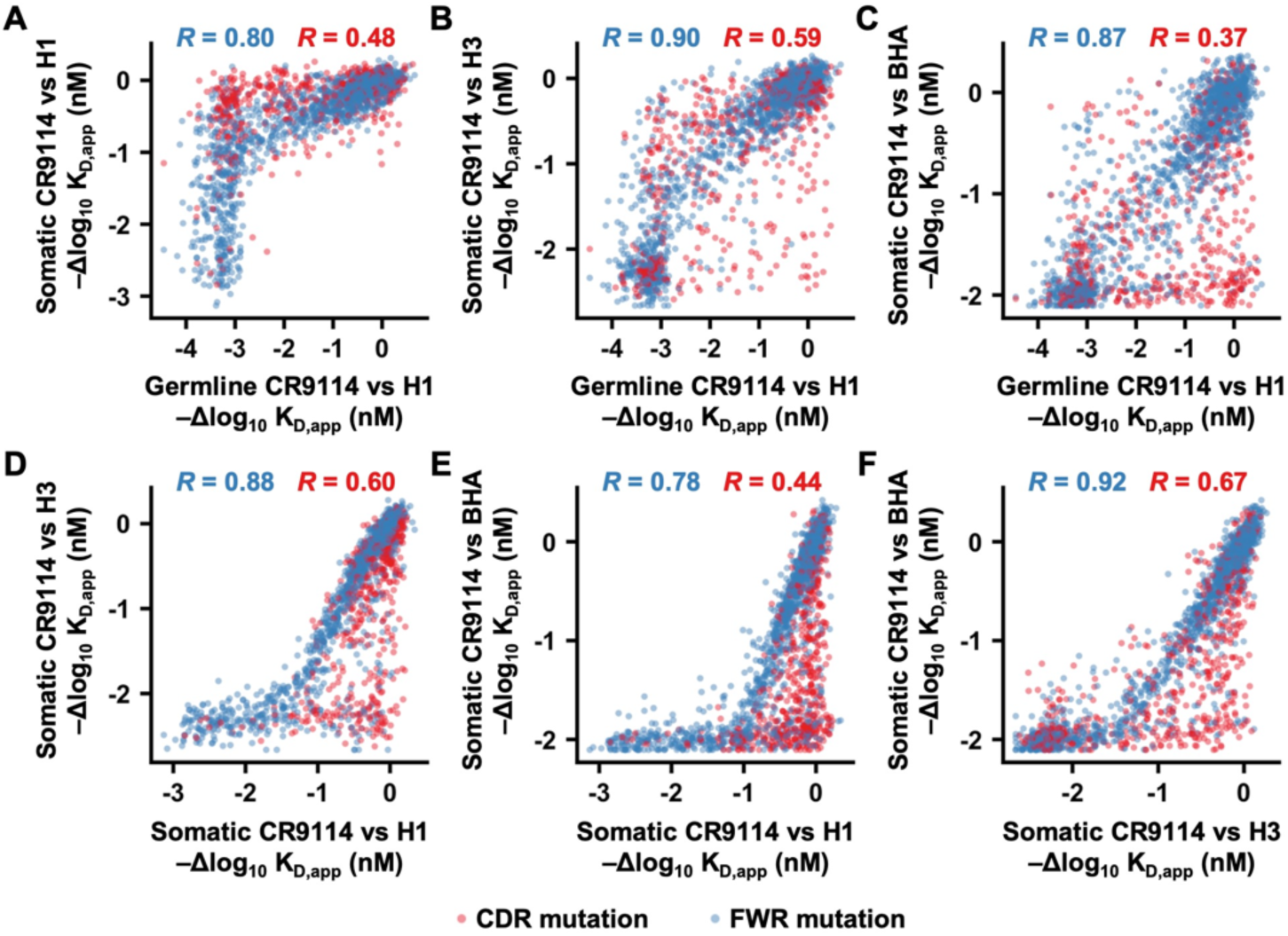
Correlation of mutational effects on binding affinity across different deep mutational scanning experiments. **(A-F)** The -Δlog_10_ K_D,app_ value of each mutation was compared between the deep mutational scanning experiments of **(A)** the germline CR9114 against H1 stem and the somatic CR9114 against H1 stem, **(B)** the germline CR9114 against H1 stem and the somatic CR9114 against H3/Sing16 HA, **(C)** the germline CR9114 against H1 stem and the somatic CR9114 against B/Lee40 HA, **(D)** the somatic CR9114 against H1 stem and the somatic CR9114 against H3/Sing16 HA, **(E)** the somatic CR9114 against H1 stem and the somatic CR9114 against B/Lee40 HA, **(F)** the somatic CR9114 against H3/Sing16 HA and the somatic CR9114 against B/Lee40 HA. Each data point represents a mutation. Pearson correlation coefficients for the mutations in the CDRs and the DE loop (red) as well as for those in the framework regions (FWRs, blue) are indicated. “H1”, “H3”, and “BHA” indicate H1 stem, H3/Sing16 HA, and B/Lee40 HA, respectively.

By comparing the binding affinity landscapes of somatic CR9114 against different HAs, we observed a pattern of nested constraint, similar to that described by a previous study on 16 somatic hypermutations of CR9114^16^. For example, while somatic CR9114 mutations that were detrimental for H1 stem binding were also detrimental for H3/Sing16 HA and B/Lee40 HA binding, many somatic CR9114 mutations that impaired H3/Sing16 HA and B/Lee40 HA binding had minimal effects on H1 stem binding (**Figure 3D-E and Figure S7D-E**). Additionally, while somatic CR9114 mutations that were detrimental for H3/Sing16 HA binding were also detrimental for binding to B/Lee40 HA, many somatic CR9114 mutations that impaired B/Lee40 HA binding had minimal effects on H3/Sing16 HA binding (**Figure 3F and Figure S7F**). Notably, most mutations that exhibited differential effects on binding against different HAs were in the CDRs (**Figure 3D-F and Figure S7D-F**). Together, our results suggested that further improving the affinity of somatic CR9114 against H1 HA could result in a loss of cross-reactive breadth due to pleiotropy.

### Epistasis and pleiotropy involve non-paratope mutations

To experimentally validate mutations that exhibited differential effects on the binding affinity of the germline and somatic CR9114 against different HAs, we recombinantly expressed and purified mutants of interest, then measured their binding affinity using biolayer interferometry (**Figure S3 and Figure S4**). The -Δlog_10_ K_D,app_ values from Tite-Seq had a Pearson correlation coefficient of 0.87 with the -Δlog_10_ K_D_ values from biolayer interferometry (**Figure 4A**). Mutations in this validation experiment included V_H_ S24F and V_H_ S24L, which abolished the binding of somatic CR9114 to B/Lee40 HA, but not to H1 stem and H3/Sing16 HA (**Figure 4B, Figure S3, and Figure S4**). Our biolayer interferometry results also showed that V_H_ I52G, V_H_ I52V, and V_H_ K73W had minimal effects on the binding of germline CR9114 to the H1 stem, whereas V_H_ S52G, V_H_ S52V, and V_H_ I73W slightly weakened the binding of somatic CR9114 to H1 stem while completely abolishing its binding to H3/Sing16 HA and B/Lee40 HA (**Figure 4B, Figure S3, and Figure S4**). Our validation experiment substantiated that epistasis and pleiotropy could constrain the breadth evolution of CR9114.

**Figure 4.**
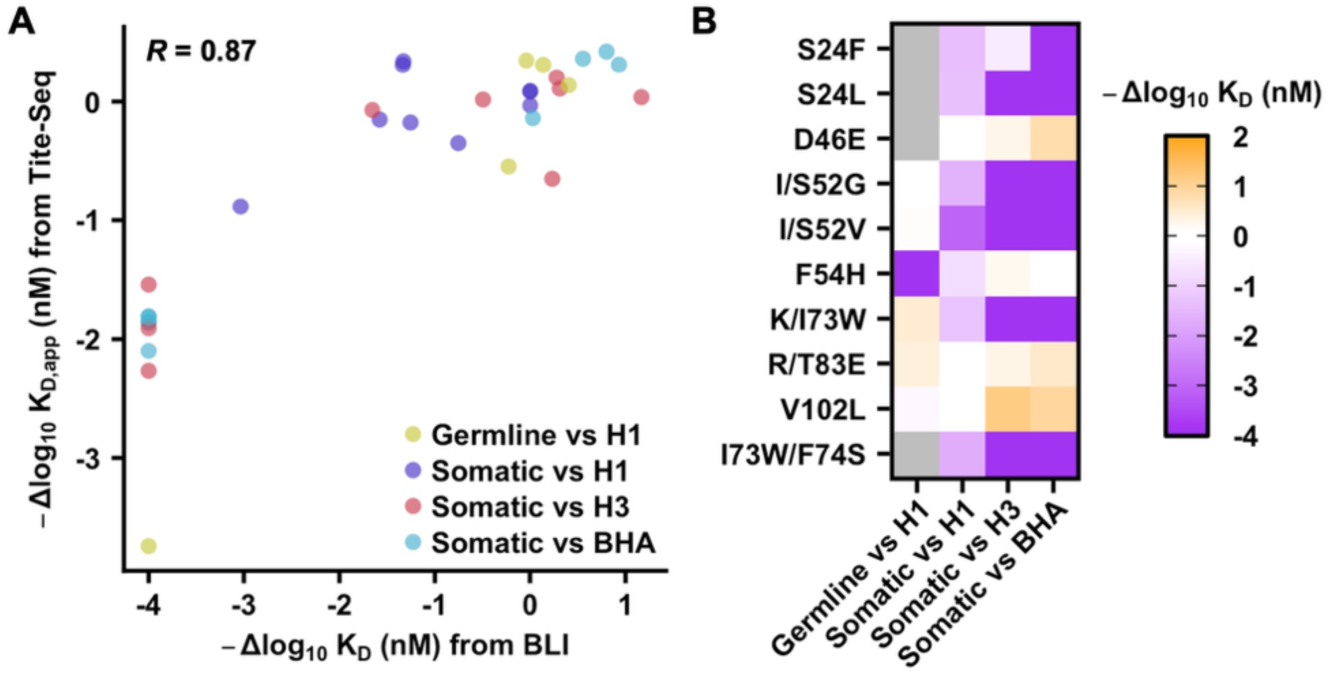
Experimental validation of the deep mutational scanning results. **(A)** The -Δlog_10_ K_D_ values determined by biolayer interferometry (BLI) are plotted against the -Δlog_10_ K_D,app_ values determined by Tite-Seq in the deep mutational scanning experiments. Each data point represents a mutation. Mutations with no detectable binding in BLI are assigned a -Δlog_10_ K_D_ value of -4. The genetic background of each mutation (i.e. germline or somatic CR9114) and the binding target (i.e. H1 stem, H3/Sing16 HA, or B/Lee40 HA) are color coded. Pearson correlation coefficient (R) is indicated. **(B)** Heatmap of BLI results for the indicated mutant in the indicated genetic background (i.e. germline or somatic variant CR9114) against the indicated target (i.e. H1 stem, H3/Sing16 HA, or B/Lee40 HA). “H1”, “H3”, and “BHA” indicate H1 stem, H3/Sing16 HA, and B/Lee40 HA, respectively.

While V_H_ residue 73 is part of the CR9114 paratope (i.e. antibody region that contacts the epitope), V_H_ residues 24 and 52 are not (**Figure 5A**). In the somatic CR9114, V_H_ S24 H-bonds with the backbone carbonyl of V_H_ G27 and the side chain of V_H_ S29 to stabilize the conformation of CDR H1 (**Figure 5B**). Mutations V_H_ S24F and V_H_ S24L would abolish these H-bonds and increase the conformational flexibility of CDR H1. Together with our biolayer interferometry results (**Figure 4B, Figure S3, and Figure S4**), this observation suggested that the conformational stability of CDR H1 was required for somatic CR9114 to bind to BHA, but not to H1 HA and H3 HA. As for V_H_ residue 52, the germline CR9114 had an Ile, whereas the somatic CR9114 had a Ser. In the somatic CR9114, V_H_ S52 H-bonds with the backbone carbonyl of V_H_ Y98 to stabilize the conformation of both CDR H2 and CDR H3 (**Figure 5C**). However, this H-bond between CDR H2 and CDR H3 would be absent in the germline CR9114 due to its hydrophobic V_H_ I52. As a result, while mutating V_H_ residue 52 in the somatic CR9114 to Gly or Val would abolish an intramolecular H-bond, the same would not occur in the germline CR9114. This observation could explain the differential effects of V_H_ I/S52G and V_H_ I/S52V on the binding affinity of somatic and germline CR9114 (**Figure 4B, Figure S3, and Figure S4**).

**Figure 5.**
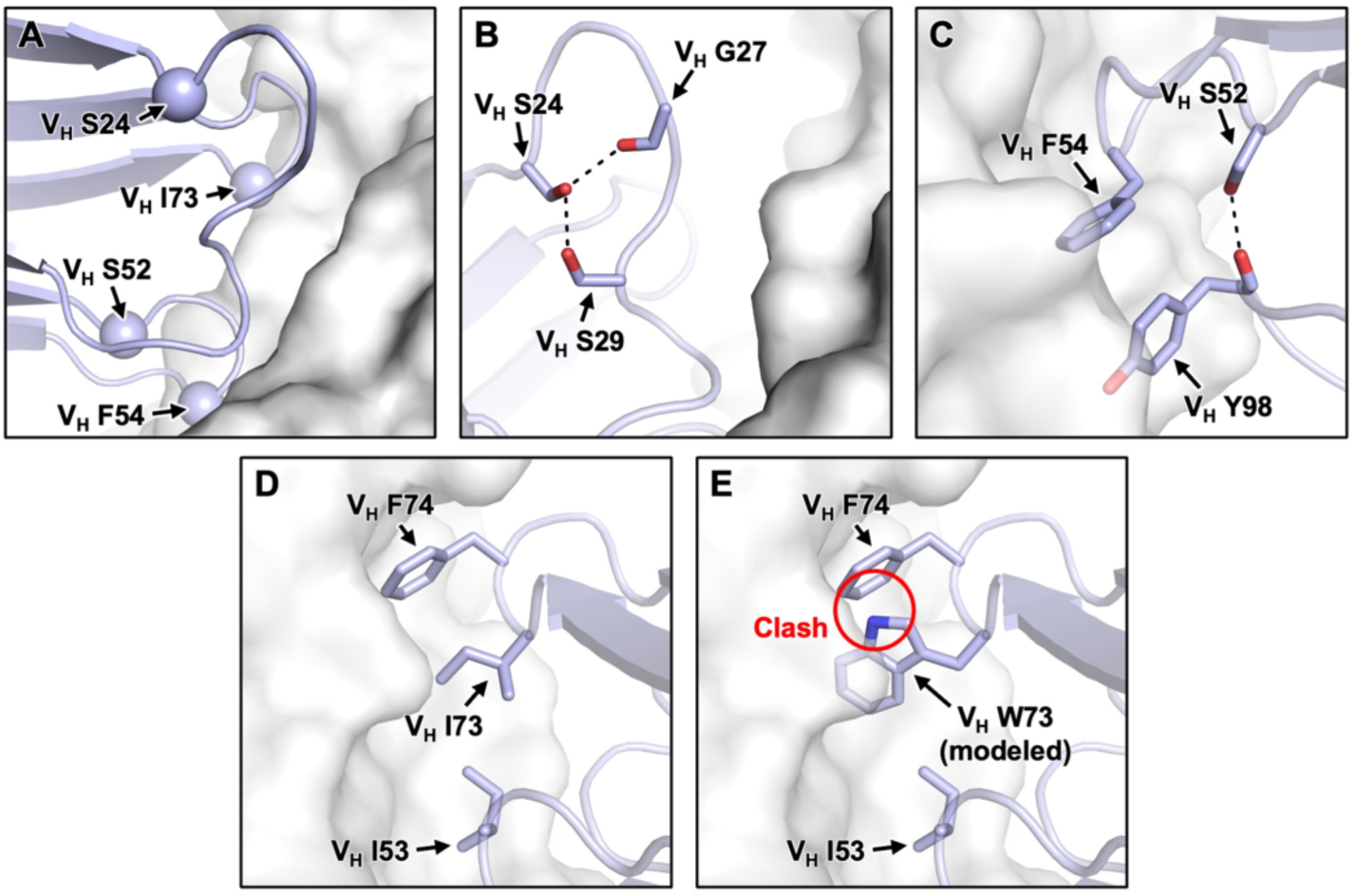
Structural analysis of the differential effects on binding affinity. **(A)** The Cαs of V_H_ S24, V_H_ S52, V_H_ F54, and V_H_ I73 in CR9114 are shown as spheres (PDB 4FQI)^15^. CR9114 and HA are in blue and white, respectively. The local chemical environment for **(B)** V_H_ S24, **(C)** V_H_ S52 and V_H_ F54, and **(D)** V_H_ I73 are shown. Black dashed lines represent H-bonds. **(E)** The structure of HA-bound CR9114 with V_H_ W73 is modeled. Red circle indicates steric clash.

### A paratope mutation constrains the breadth evolution of CR9114

Our biolayer interferometry results showed that mutating V_H_ residue 73 to Trp improved binding of germline CR9114 H1 stem, but abolished the binding of somatic CR9114 to H3/Sing16 HA and B/Lee40 HA (**Figure 4B, Figure S3, and Figure S4**). V_H_ residue 73 is part of the CR9114 paratope. In somatic CR9114, V_H_ I73 is adjacent to V_H_ F74 (**Figure 5D**). Our structural modelling indicated that mutation V_H_ I73W would introduce a steric clash with V_H_ F74, which could explain its detrimental effect on the binding of somatic CR9114 to H3/Sing16 HA and B/Lee40 HA. By contrast, this detrimental steric clash was not observed in the germline CR9114, which has the small amino acid Ser at V_H_ residue 74 (**Figure 2**). However, somatic CR9114 with double mutations V_H_ I73W/F74S was still unable to bind to H3/Sing16 HA or B/Lee40 HA (**Figure 4B and Figure S4**). Therefore, V_H_ W73 could impose strong constraints for CR9114 to evolve breadth, which may not be easily overcome by a secondary mutation.

### Epistasis is prevalent in IGHV1-69 HA stem antibodies

CR9114 is encoded by IGHV1-69, which is a recurring sequencing feature of HA stem antibodies^21–23^. Our previous study has curated over 5000 influenza HA antibodies from the literature, of which 160 are IGHV1-69 HA stem antibodies^24^. Here, we analyzed the somatic hypermutations present in these 160 IGHV1-69 HA stem antibodies in terms of their effects on the binding affinity of germline CR9114. Most of the IGHV1-69 HA stem antibodies contained at least one somatic hypermutation with a -Δlog_10_ K_D,app_ value of less than -1 (**Figure 6**), which was a stringent cutoff for deleterious mutations according to our validation experiment (**Figure 4A**). This observation demonstrated the prevalence of epistasis among the affinity maturation trajectories of IGHV1-69 HA stem antibodies. This epistasis could be due to a given mutation having different effects on the binding affinity of the germline precursors of different IGHV1-69 HA stem antibodies. Consistently, different binding modes have been reported for IGHV1-69 HA stem antibodies^22,25^. Another possibility is that a given mutation may be detrimental to the germline precursors of all IGHV1-69 HA stem antibodies, but may become neutral or beneficial in the presence of other somatic hypermutations, as observed in CR9114^16^. Regardless, our results suggested that epistasis is prevalent among the binding affinity landscapes of IGHV1-69 HA stem antibodies and may constrain their evolution of breadth.

**Figure 6.**
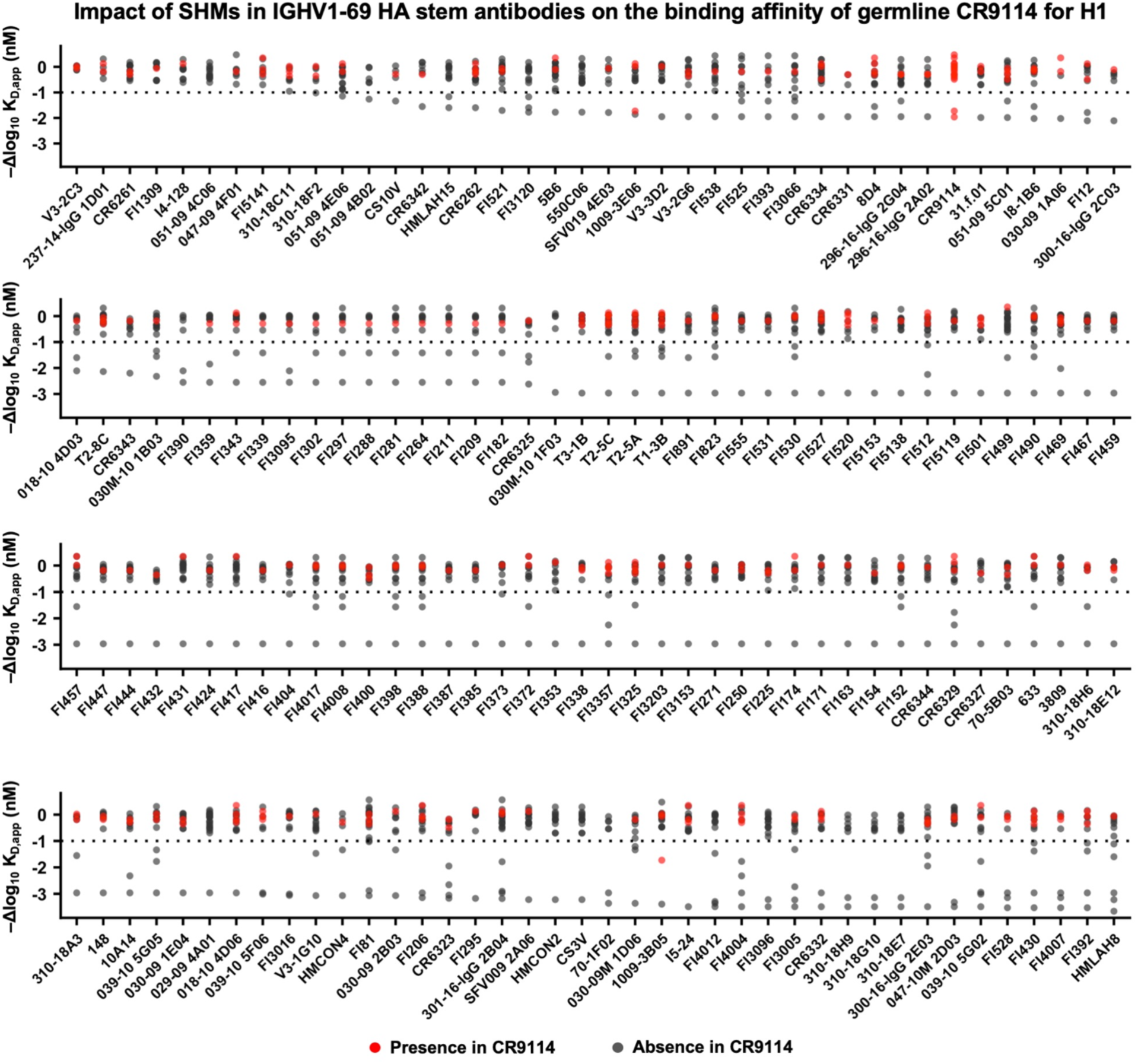
The effects of somatic hypermutations in IGHV1-69 HA stem antibodies on the binding affinity of germline CR9114. The somatic hypermutations in each of the 160 known IGHV1-69 HA stem antibodies^24^ were identified. Their corresponding -Δlog_10_ K_D,app_ values obtained from the deep mutational scanning experiment of the germline CR9114 against H1 stem (Figure 2) are shown. Each data point represents one somatic hypermutation of the indicated antibody. Somatic hypermutations observed in CR9114 are in red. The name of each antibody is shown on the x-axis.

## DISCUSSION

CR9114 was one of the first few bnAbs discovered against influenza HA stem^15^. Given that CR9114 is encoded by the commonly used IGHV1-69 germline gene and lacks unusual features that are observed in HIV bnAbs, such as extensive somatic hypermutation^26^, long indels^27^, or domain swapping^28^, CR9114-like antibodies were initially thought to be readily generable and present in different individuals^15^. However, despite being discovered more than a decade ago, CR9114 remains the only bnAb found to date that cross-reacts with both influenza A and B HAs. In fact, most IGHV1-69 HA stem antibodies are specific to group 1 HA and cannot even cross-react with group 2 HA^24,29^. To understand the potential evolutionary constraints that would limit the elicitation of CR9114-like antibodies, this study systematically analyzed the mutational effects on the binding affinity of the germline and somatic CR9114 against multiple HAs. We found that the breadth evolution of CR9114 is highly vulnerable due to epistasis and pleiotropy. Our results not only help explain the rarity of HA stem bnAbs that cross-react with influenza A and B viruses, but also have important implications for developing broadly protective influenza vaccines.

A notable finding in this study was that the mutational tolerance of somatic CR9114 decreased in correlation with its binding affinity, which weakened sequentially across the H1 stem, H3 HA, and BHA. This observation is substantiated by our validation experiment, where mutations V_H_ S52G, V_H_ S52V, and V_H_ I73W abolished the binding of somatic CR9114 to H3 HA and BHA, but not to H1 stem. Consistently, the mutational tolerance and binding affinity of germline CR9114 against H1 stem were similar to those of somatic CR9114 against BHA. This phenomenon parallels a previous study which showed the presence of sequence determinants that are functionally redundant for H1 HA binding in somatic but not germline IGHV1-69 HA stem antibodies^21^. However, the underlying biophysical mechanisms and its generalizability to other classes of antibodies warrant additional studies.

Phillips et al. have previously shown that the breadth evolution of CR9114 follows a sequential gain-of-affinity in the order of H1 HA, H3 HA, then BHA, due to nested evolutionary constraints also observed in our study here^16^. Nevertheless, a sequential vaccination scheme is unlikely to be sufficient for reliably eliciting CR9114-like bnAbs, due to the large number of potential mutations that are detrimental to H3 HA and BHA binding without affecting H1 HA binding. As a result, even if CR9114-like bnAb precursors can be primed by H1 HA, they have a high chance of acquiring somatic hypermutations that prevent the subsequent acquisition of cross-reactivity breadth, especially since selection in the germinal centers is weak^30^. Compared to the difficulties in eliciting CR9114-like bnAbs that cross-react with influenza A and B viruses, the generation of HA stem bnAbs with pan-influenza A reactivity appeared much easier^31–33^. For example, a single amino acid mutation is able to confer cross-group reactivity to the multidonor HA stem bnAbs encoded by IGHV1-18 with a QxxV motif in the CDR H3^31^. Therefore, we postulate that a reasonable strategy for developing broadly protective influenza vaccines against both human and zoonotic strains is to use one immunogen to elicit pan-influenza A bnAbs and a second immunogen for pan-influenza B bnAbs.

To date, multiple HA-stem based influenza vaccine candidates have been developed^12,18,34–37^. Nevertheless, both group 1 and group 2 HA stem-based vaccine candidates tested in clinical studies have shown limited ability to elicit cross-group bnAbs, despite robustly eliciting group-specific bnAbs^38–41^. The lack of a reliable method for eliciting cross-group HA stem bnAbs represents a major obstacle toward a pan-influenza A vaccine. At the same time, a recent study has demonstrated that the antibody evolutionary outcomes in germinal centers are predictable based on the antibody binding affinity landscapes and somatic hypermutation biases^30^. Therefore, continued characterization of the binding affinity landscapes of diverse HA stem bnAbs will be crucial for guiding the rational development of a pan-influenza A vaccine.

## ACKNOWLEDGEMENTS

This work is supported by National Institutes of Health R01 AI167910 (N.C.W.), DP2 AT011966 (N.C.W.), and the Vallee Scholars Program (N.C.W.). We thank the Roy J. Carver Biotechnology Center at the University of Illinois at Urbana-Champaign for assistance with cell sorting and next-generation sequencing.

## AUTHOR CONTRIBUTIONS

N.C.W. and K.E.D. conceived and designed the study. K.E.D. performed the experiments. Y.W. and C.W. performed the data analysis. Q.W.T. provided technical expertise in Tite-Seq. R.L., M.T., L.A.R., and L.W. expressed and purified the recombinant HA proteins. K.E.D. and N.C.W. wrote the paper, and all authors reviewed and edited the paper. Thanks to Tossapol Pholcharee and Wenhao Owen Ouyang for helpful discussions.

## METHODS

### Yeast display plasmids

Yeast display plasmids for germline and somatic CR9114 Fabs were constructed based on the pCTcon2 vector, with an open reading frame encoding (from N-terminal to C-terminal): Aga2 secretion signal, somatic CR9114 Fab light chain, V5 tag, equine rhinitis B virus (ERBV-1) 2A peptide, Aga2 secretion signal, germline or somatic CR9114 Fab heavy chain, HA-tag, and Aga2p.

### Construction of yeast Fab mutant libraries

Two separate mutant libraries were generated based on different versions of the CR9114 heavy chain variable domain sequence. The fully developed or somatic CR9114 sequence, and the pre-developmental, or germline CR9114 sequence, were each used as bases for the respective libraries. The heavy and light chain sequences of somatic CR9114 were obtained from Genbank JX213639 and JX213640, respectively (*1*). The CR9114 germline sequence was obtained from Phillips et al. (*2*), who themselves had reconstructed the sequence through IMGT (*3*) and IgBLAST (*4*). We chose to mimic their treatment of site S109N, and not acknowledge a mutation at that site (*2*). For both libraries, the somatic CR9114 light chain sequence was used, to focus solely on heavy chain variant effects. The mutant libraries of variant Fabs were generated that differed from the somatic or germline CR9114 sequences by only one amino acid per mutant, such that every possible amino acid, as well as STOP codons, was represented in every possible position of the heavy chain variable domain, excluding the final amino acid in the heavy chain variable domain, which is highly conserved.

The pCTcon2 plasmid containing either the somatic or germline CR9114 sequence in Fab form was used as a template to generate the respective mutant library inserts and the linearized vector through polymerase chain reaction (PCR), separately. The inserts for each library were then generated as follows, per library. To generate the linearized vectors, we used 5′-TGG TGT CCA CAC TTT CCC TGC TGT T-3′ and 5′-AGT GGA TTG GGG ATT GGC TTT CCG C-3′ as primers.

To generate the full-length inserts, we first needed to manufacture two sets of 15 split inserts per library in parallel before merging them in paired PCRs using overlap extension PCR. The first set of split inserts included 15 separate reactions; each performed with a cassette of pooled forward primers and a universal reverse primer 5′-AGG AGT ACA AAC CGG AAG ATT GC-3′. Each forward cassette of primers was composed of eight forward primers in equal molar ratios with an identical 21 nt at the 5’ end and 15 nt at the 3’ end. NNK (N: A, C, G, T; K: G, T) sequences were present within each cassette, placed in such a way that saturation mutagenesis was achieved.

As described previously, silent (or synonymous) mutations were introduced into each primer to discern between sequencing errors and target mutations through data analysis (*5*). In **Table S2**, the forward primers are listed for each mutant library, identified in the table by cassette number and the position of the amino acid mutated. The second set of split inserts also included 15 separate reactions, designed to match up with the first set of split inserts. This final set of 15 insert fragments was generated through PCR, with each reaction using the universal forward primer 5′-ACT GTC GCC CCA ACT GAG TGC TC-3′ and a unique reverse primer (**Table S3**). The first and second sets of split inserts were joined to generate full-length cassettes through 15 overlapping PCRs, using universal forward and reverse primers to perform the amplifications with 10 ng of the respective inserts per PCR. For each library, these 15 full-length cassettes were gel purified, then mixed using equimolar amounts of each cassette to generate the final insert pool for use in the yeast transformation. PrimeSTAR Max polymerase (Takara Bio, cat no. R045B) was used for all PCRs. Gel purification was performed using the Monarch Gel Extraction Kit (New England Biolabs, cat no. T1020L). The manufacturer’s instructions were used for all samples in each case.

### Yeast transformation

Transformation of yeast was performed for each mutant library by electroporation as previously described (*6*). Strain EBY100 of *Saccharomyces cerevisiae* (ATCC) were grown overnight at 30C with 250 rpm agitation in YPD medium (1% w/v yeast nitrogen base, 2% w/v peptone, 2% w/v D(+)-glucose). Once the culture reached OD_600_ of 3, a new culture was seeded at OD_600_ of 0.3 in 100 mL of YPD media and continued to grow at 30°C with 250 rpm agitation. Following OD_600_ reaching 1.6, the culture of yeast cells was centrifuged for 3 min at 1700×*g* at room temperature, and the media was removed. 50 mL of ice-cold water was used to wash the cell pellet twice; the pellet was then washed with 50 mL of ice-cold electroporation buffer (1 M sorbitol, 1 mM calcium chloride) before resuspension in 20 mL conditioning media (0.1 M lithium acetate, 10 mM dithiothreitol). The new mixture was then shaken at 250 rpm at 30°C for 30 min, then centrifuged at 1700×*g* for 3 min at room temperature. Following wash with 50 mL ice-cold electroporation buffer, the pellet was resuspended in 1 mL of electroporation buffer and placed on ice. In parallel, 4 µg of the linearized vector was mixed with 5 µg of the full-length cassette mutant library insert, and pipetted into 400 µL of the prepared 1 mL conditioned yeast in a pre-chilled BioRad GenePulser 2 mm electrode gap cuvette. Following a 5-min incubation on ice, electroporation was performed at 2.5 kV, 25 µF, such that the time constant was within 3.7 and 4.1 ms. The electroporated cell mixture was immediately transferred into 8 mL of a 1 to 1 mixture of YPD and 1 M sorbitol, then shaken at 30°C, 250 rpm for 1 h.

Cells were collected via centrifugation at 1700×*g* for 3 min at room temperature, resuspended in 0.6 mL SD-CAA medium (2% w/v D-glucose, 0.67% w/v yeast nitrogen base with ammonium sulfate, 0.5% w/v casamino acids, 0.54% w/v Na_2_HPO_4_, 0.86% w/v NaH_2_PO_4_•H_2_O, all dissolved in deionized water), plated onto SD-CAA plates (2% w/v D-glucose, 0.67% w/v yeast nitrogen base with ammonium sulfate, 0.5% w/v casamino acids, 0.54% w/v Na_2_HPO_4_, 0.86% w/v NaH_2_PO_4_•H_2_O, 18.2% w/v sorbitol, 1.5% w/v agar, all dissolved in deionized water) and incubated at 30°C for 48 h. Colonies were then collected in SD-CAA medium, centrifuged at 1700×*g* for 5 min at room temperature, and resuspended in SD-CAA medium with 15% v/v glycerol such that OD600 was 50. Glycerol stocks were stored at -80°C until used.

### Cell Culture

Sf9 cells (*Spodoptera frugiperda* ovarian cells, female, ATCC, cat no. CRL-1711) were maintained in Sf-900 II SFM medium (Thermo Fisher Scientific, cat no. 10902088) at 37°C. Expi293F cells (Gibco, cat no. A14527) were grown and maintained in Expi293 Expression Medium (Gibco, cat no. A1435101) at 37°C, 8% CO_2_, and 95% humidity with shaking at 125 rpm according to the manufacturer’s instructions.

### Expression and purification of H1 stem, H3 HA, and BHA

Oligonucleotide encoding H1 stem #4900 protein (*7*), H3 HA (A/Singapore/INFIMH-16-0019/2016; GISAID accession num. EPI1858150), or BHA (B/Lee/1940; GISAID accession num. EPI243230) was fused with N-terminal gp67 signal peptide and a C-terminal BirA biotinylation site, thrombin cleavage site, trimerization domain, and a 6× His tag, and then cloned into a customized baculovirus transfer vector (*8*). Recombinant bacmid DNA that carried the protein construct was generated using the Bac-to-Bac system (Thermo Fisher Scientific, cat no. 10359016) according to the manufacturer’s instructions. Baculovirus was generated by transfecting the purified bacmid DNA into adherent Sf9 cells using Cellfectin reagent (Thermo Fisher Scientific, cat no. 10362100) according to the manufacturer’s instructions. The baculovirus was further amplified by passaging in adherent Sf9 cells at a multiplicity of infection (MOI) of 1. Recombinant protein was expressed by infecting 1 L of suspension Sf9 cells at an MOI of 1. Infected Sf9 cells were incubated at 27°C with shaking at 170 rpm for 72 h. On day 3 post-infection, Sf9 cells were pelleted by centrifugation at 4000×*g* for 25 min, and soluble recombinant protein was purified from the supernatant by affinity chromatography using Ni Sepharose excel resin (Cytiva, cat no. 17371202) and then size exclusion chromatography using a HiLoad 16/100 Superdex 200 prep grade column (Cytiva, cat no. 90100137) in 20 mM Tris-HCl pH 8.0, 100 mM NaCl. The purified protein was concentrated by Amicon spin filter (Millipore Sigma, cat no. UFC901008) and filtered by 0.22 mm centrifuge Tube Filters (Costar, cat no. 8160). Concentration of the protein was determined by a NanoDrop One^C^ spectrophotometer (Fisher Scientific). Protein was subsequently aliquoted, flash frozen in dry-ice ethanol mixture, and stored at -80°C until use.

### Biotinylation and PE-conjugation of HAs

Purified H1 stem, H3 HA, and BHA were biotinylated using the Biotin-Protein Ligase-BIRA kit according to the manufacturer’s instructions (Avidity, cat no. BirA500). After buffer exchange into PBS, the biotinylated HA were conjugated to streptavidin-PE (Fisher Scientific, cat no. S866A) by incubating at room temperature for 15 min.

### Fluorescence-activated cell sorting (FACS) of yeast display libraries

100 µL glycerol stock of the yeast display library was recovered in 50 mL SD-CAA medium by incubating at 27°C with shaking at 250 rpm until OD_600_ reached between 1.5 and 2.0. Then 15 mL of the yeast culture was harvested and pelleted via centrifugation at 4000×*g* at 4°C for 5 min. The supernatant was discarded, and SGR-CAA (2% w/v galactose, 2% w/v raffinose, 0.1% w/v D-glucose, 0.67% w/v yeast nitrogen base with ammonium sulfate, 0.5% w/v casamino acids, 0.54% w/v Na_2_HPO_4_, 0.86% w/v NaH_2_PO_4_•H_2_O, all dissolved in deionized water) was added to make up the volume to 50 mL. The yeast culture was then transferred to a baffled flask and incubated at 18°C with shaking at 250 rpm. Once OD_600_ reached between 1.3 and 1.6, 1 mL of yeast culture was harvested and pelleted via centrifugation at 4000×*g* at 4°C for 5 min. The pellet was subsequently washed with 1 mL of 1× PBS twice. After the final wash, cells were resuspended in 1 mL of 1× PBS. For each expression sort, PE anti-HA.11 (epitope 16B12, BioLegend, cat. no. 901517) that had been buffer-exchanged into 1× PBS was added to the cells at a final concentration of 30 nM.

For the expression sorts, PE anti-HA.11 (Biolegend, cat no. 901518) was added to a subset of the washed cells at a final concentration of 30 nM. For Tite-Seq, cells were labeled with PE-conjugated H1 stem, H3 HA, or BHA at each of eight antigen concentrations. For the somatic CR9114 mutant library sorts, one-log increments of PE-conjugated probe spanning 0.01 nM–10 nM and 0.003 nM–3 nM (H1 stem); 0.1 nM–100 nM and 0.03 nM–33 nM (H3 HA); or 0.2 nM–200 nM and 0.067 nM–67 nM (BHA) were used for FACS. For the germline CR9114 mutant library sorts, one-log increments of PE-conjugated H1 stem spanning 16 nM–0.016 nM and 5 nM–0.005 nM, along with an unstained sample, were used for FACS. A negative control was set up with no probe added to the resuspended cells. For the somatic CR9114 mutant library sorts, all cells were stained in solution containing 1× PBS with 1% BSA, while for the germline CR9114 mutant library sorts, all cells were stained in solution of 1× PBS lacking BSA. The reason for this discrepancy is that the method was improved during this project: the addition of 1% BSA was found to provide a cleaner sort with lower background signal. Samples were incubated with their respective probe concentrations for 1 h at 4°C with rotation. Then, the yeast pellet was washed twice in 1× PBS and resuspended in FACS tubes containing 2 mL 1× PBS. Cells in the selected gates were collected in 1 mL of SD-CAA containing 1× penicillin/streptomycin. The somatic CR9114 mutant library was sorted using a BD FACSMelody Cell Sorter and BD FACSChorus Software (BD Biosciences), while the germline CR9114 mutant library was sorted using a BD FACSAria and BD FACSDiva Software (BD Biosciences).

For expression sort, single cells were gated into four equally populated bins, with bin 1 to 4 having the increasing PE signal. For Tite-Seq, single cells were gated into four bins along the PE-A axis based on unstained and CR9114 controls, with bin 1 comprising all PE negative cells, bin 4 comprising PE positive cells with comparable expression or binding affinity to the germline or somatic CR9114 positive population, and bins 2 and 3 splitting the intermediate population between bin 1 and bin 4. Following sort, the sort tubes were directly placed at 30°C with 250 rpm agitation. After 20-40 h, the cells in each tube were collected by centrifugation at 3800×*g* for 15 min once a large pellet was visible or the culture was cloudy. Frozen stocks were made per sorted sample by reconstituting the pellet in 15% v/v glycerol (in SD-CAA medium) and then stored at - 80°C until use.

### Next-generation sequencing of FACS samples

Plasmids from the yeast cells before and after sorting were extracted using a Zymoprep Yeast Plasmid Miniprep II Kit (Zymo Research, cat no. D2004) following the manufacturer’s protocol. The region of interest in the mutant library was subsequently amplified by PCR using primers 5’-C ACT CTT TCC CTA CAC GAC GCT CTT CCG ATC TGT GTA CTT GCC GCC GCT CAA CCA -3’ and 5’-G ACT GGA GTT CAG ACG TGT GCT CTT CCG ATC TAC GGA AGG TCC CTT AGT AGA AGC-3’ for both the germline and somatic mutant libraries. Subsequently, adapters containing sequencing barcodes were appended to the amplicon using primers 5’-AAT GAT ACG GCG ACC ACC GAG ATC TAC ACX XXX XXX XAC ACT CTT TCC CTA CAC GAC GCT-3’, and 5’-CAA GCA GAA GAC GGC ATA CGA GAT XXX XXX XXG TGA CTG GAG TTC AGA CGT GTG CT-3’. Positions annotated by an ‘‘X’’ represented the nucleotides for the index sequence. All PCRs post-FACS were performed using Q5 High-Fidelity DNA polymerase (NEB, cat no. M0492S) according to the manufacturer’s instructions. PCR products were purified using PureLink PCR Purification Kit (Thermo Fisher Scientific, cat no. K310002). The final PCR products were submitted for next generation sequencing using NovaSeq SP PE250 (Illumina).

### Computing the binding score, expression score, and apparent K_D_ values

The sequencing data were initially obtained in FASTQ format, then analyzed using a custom python Snakemake pipeline^42^. Briefly, PEAR was used to merge the forward and reverse reads^43^. The number of reads corresponding to each variant in each sample was counted. To avoid division by zero in downstream analysis, a pseudocount of 1 was added to the final count.

For each expression sort, the enrichment value of each mutation *mut* were computed as follows:

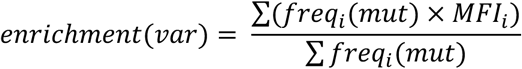

where the *freq_i_*(*var*) is the occurrence frequency of mutation *mut* in bin *i* and *MFI_i_* is the mean fluorescence intensity of bin *i*. The expression score for each variant *var* was further computed from the enrichment value as follows:

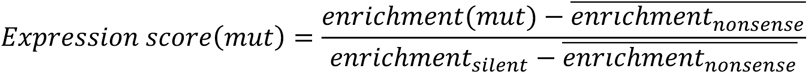

where 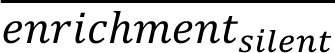 is the average enrichment value of the silent mutations and 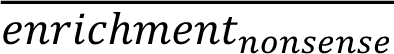 is the average enrichment value of the nonsense mutations. The final score for each variant *var* is the average of two biological replicates.

To compute apparent *K_D_* value of each variant from the Tite-Seq data, we adopted the analysis approach as previously described^44^. Briefly, to determine the mean bin of PE fluorescence for each variant *var* at each HA concentration, a simple weighted mean calculation was applied:

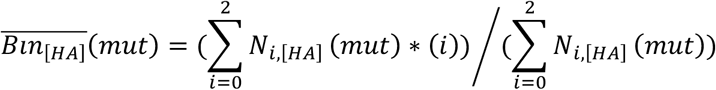

where *N*_*i*,_[*HA*](*mut*) is the number of cells with mutation *mut* that fall into bin *i* at HA concentration [*HA*]. This calculation computes a weighted average by assigning integer weight to the bin *i*.

For each variant *var*, we estimated its sorted cell count *N*_*i*,[*HA*_(*mut*) that corresponds to bin *i* at HA concentration [*HA*] as follows:

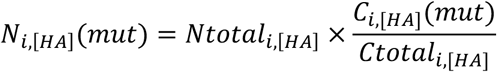

where variant read count *C*_*i*,[*HA*](*mut*) is the read counts for variant *var* in bin *i* at HA concentration [*HA*], *Ctotal_i_*,[*HA*]_ is the total read counts for bin *i* at HA concentration [*HA*], *Ntotal*_*i*,[*HA*]_ is the total number of cells in bin *i* at HA concentration [*HA*]. We then determined the apparent binding affinity *K_D,app_* value for each mutation *K_D,app_*(*mut*) via a nonlinear least-squares regression using a standard non-cooperative Hill equation:

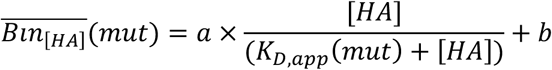

where free parameters *a* is titration response range and *b* is titration curve baseline. Mutations with an occurrence frequency below 0.01% in our pre-sort samples were discarded from our downstream analysis. To quantify the effects of each mutation on binding affinity, we calculate the difference between the *K_D,app_* values of the given mutation and the corresponding wild type:

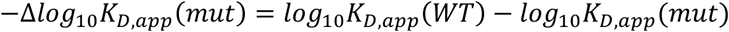

### Expression and purification of Fabs

The heavy and light chains of CR9114 Fab germline or somatic heavy chain sequence mutants were cloned into phCMV3 vector. The mutants were generated using mismatch PCR to modify the bases desired using PrimeSTAR Max polymerase (Takara Bio, catalog no. R045B). Gibson cloning was then performed using the NEBuilder HiFi DNA Assembly Master Mix (NEB, cat no. E2621), using the manufacturer’s instructions following mismatch PCR to modify the bases desired. The plasmids were co-transfected into Expi293F cells at a 2:1 (HC:LC) mass ratio using ExpiFectamine 293 Reagent (Thermo Fisher Scientific, cat no. A14525). At 5 days post-transfection, the supernatant was collected, filtered using 100.0 µm Millipore Steriflip Vacuum Tube Top Filters (Millipore Sigma, cat no. SCNY00100), and the Fab was purified using a CaptureSelect CH1-XL Affinity Matrix (Thermo Fisher Scientific, cat no. 194371201L).

### Biolayer interferometry binding assay

Binding assays were performed by biolayer interferometry (BLI) using an Octet Red96e instrument (FortéBio) at room temperature as described previously (*9*). Briefly, biotinylated H1 stem, H3 HA, or BHA proteins at 20 µg mL^-1^ in 1× kinetics buffer (PBS at pH 7.4, 0.01% w/v BSA, and 0.002% v/v Tween 20) were loaded onto streptavidin (SA) biosensors and incubated with the indicated concentrations of Fabs. The assay consisted of five steps, each run for at least 60 seconds: (1) baseline: 1× kinetics buffer; (2) loading: HA proteins; (3) baseline: 1× kinetics buffer; (4) association: Fab samples; and (5) dissociation: 1× kinetics buffer. Two to three Fab concentrations (33 nM, 100 nM and 300 nM) were used for each assay. However, for testing of S24L on the somatic CR9114 background against H3 HA, 1000, 300, and 100 nM of Fab in 1× BLI buffer were used. For estimating the K_D_, a 1:1 binding model was used. In cases where 1:1 binding model did not fit well due to the contribution of non-specific binding to the response curve, a 2:1 heterogeneous ligand model was used to improve the fitting.

### Differential scanning fluorimetry assay

Thermal shift assays were performed using differential scanning fluorimetry (DSF) on the CFX Opus 96 Real-Time PCR System (Bio-Rad, cat no. 12011319). SYPRO Orange Protein Gel Stain (5,000X Concentrate in DMSO) (Thermo Fisher Scientific, cat no. S6650). 5ug of each Fab was used per assay in a total volume of 25 µL with SYPRO Orange Protein Gel Stain at 1X in PBS in a Hard-Shell 96-Well PCR Plate (Bio-Rad, cat no. HSP9601). The assay was performed from 10°C to 95°C in increments of 0.5°C for 10 s plus plate reading using the FRET channel of the instrument. Output files were analyzed with R.

## QUANTIFICATION AND STATISTICAL ANALYSIS

Standard deviation for K_D_ estimation, as well as K_D_ itself, was computed by Octet analysis software 9.0. All statistical tests and correlation coefficients were computed in R.

## CODE AVAILABILITY

Custom python scripts for analyzing the deep mutational scanning data have been deposited to https://github.com/nicwulab/CR9114_DMS.

## DATA AVAILABILITY

Raw sequencing data have been submitted to the NIH Short Read Archive under accession number: BioProject PRJNA1284397.

