## Supplemental Figures for "Vulnerability in the breadth evolution of an influenza broadly neutralizing antibody"

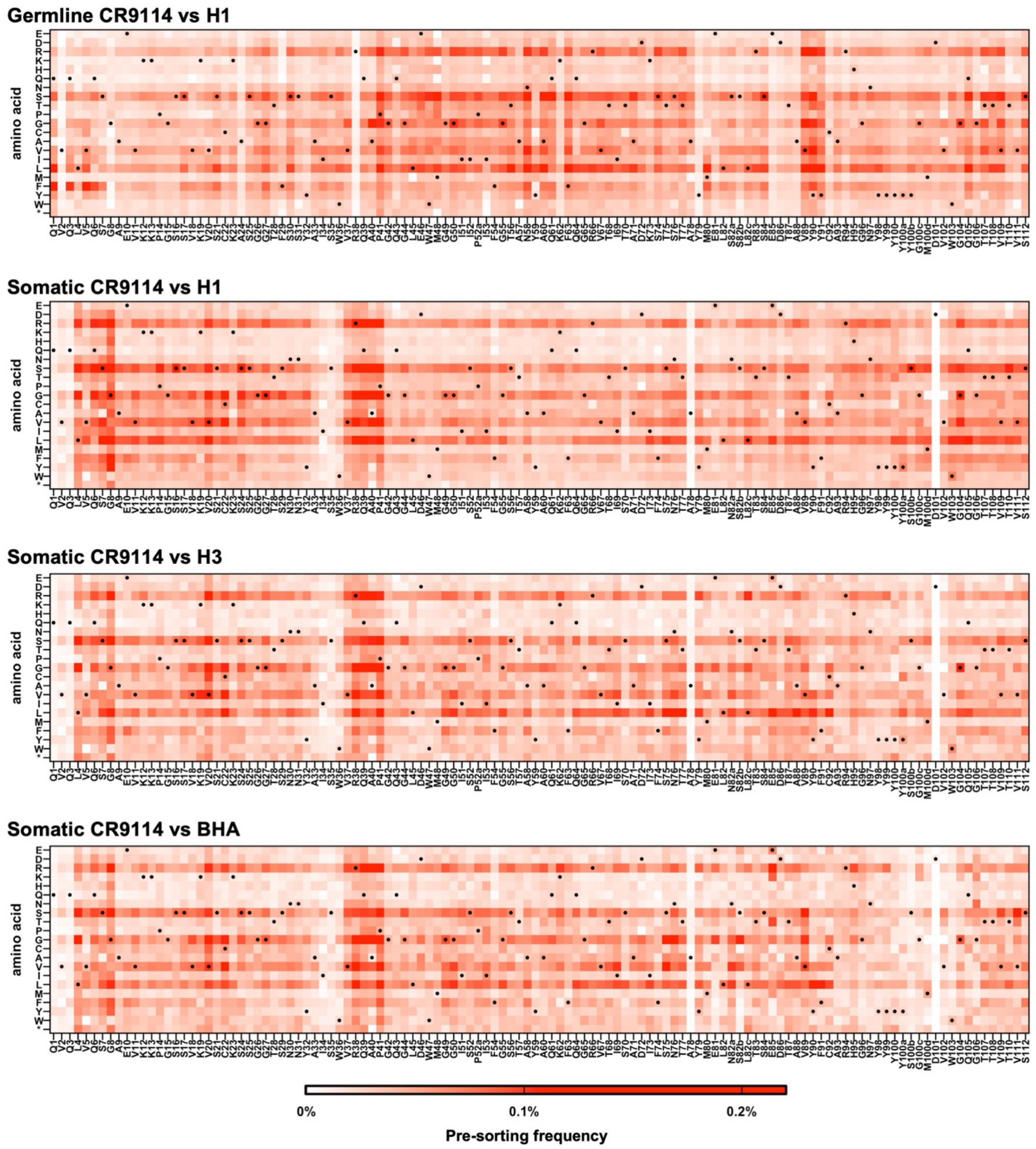


**Figure S1. Occurrence frequency of each mutation before sorting. (A-D)** Heatmaps showing the pre-sorting frequency of each mutation in each deep mutational scanning experiment. “H1”, “H3”, and “BHA” indicate H1 stem, H3/Sing16 HA, and B/Lee40 HA, respectively.

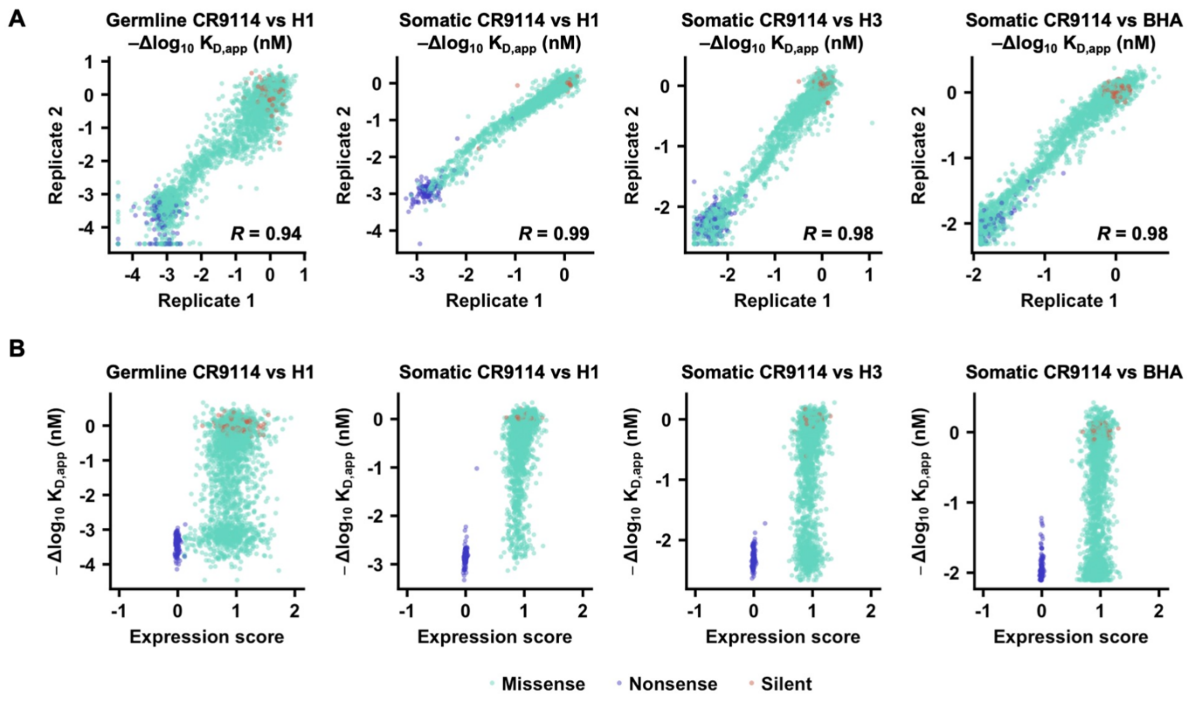


**Figure S2. Correlations between replicates and expression score comparisons during sort.** **(A)** Correlation of -$\Delta\log_{10}$ $K_{D,app}$ values between replicates **(B)** Correlation of expression scores with -$\Delta\log_{10}$ $K_{D,app}$ values. Each teal, blue, or orange dot placed at the coordinate values represents a missense, nonsense, or silent mutation, respectively. “H1”, “H3”, and “BHA” indicate H1 stem, H3/Sing16 HA, and B/Lee40 HA, respectively.

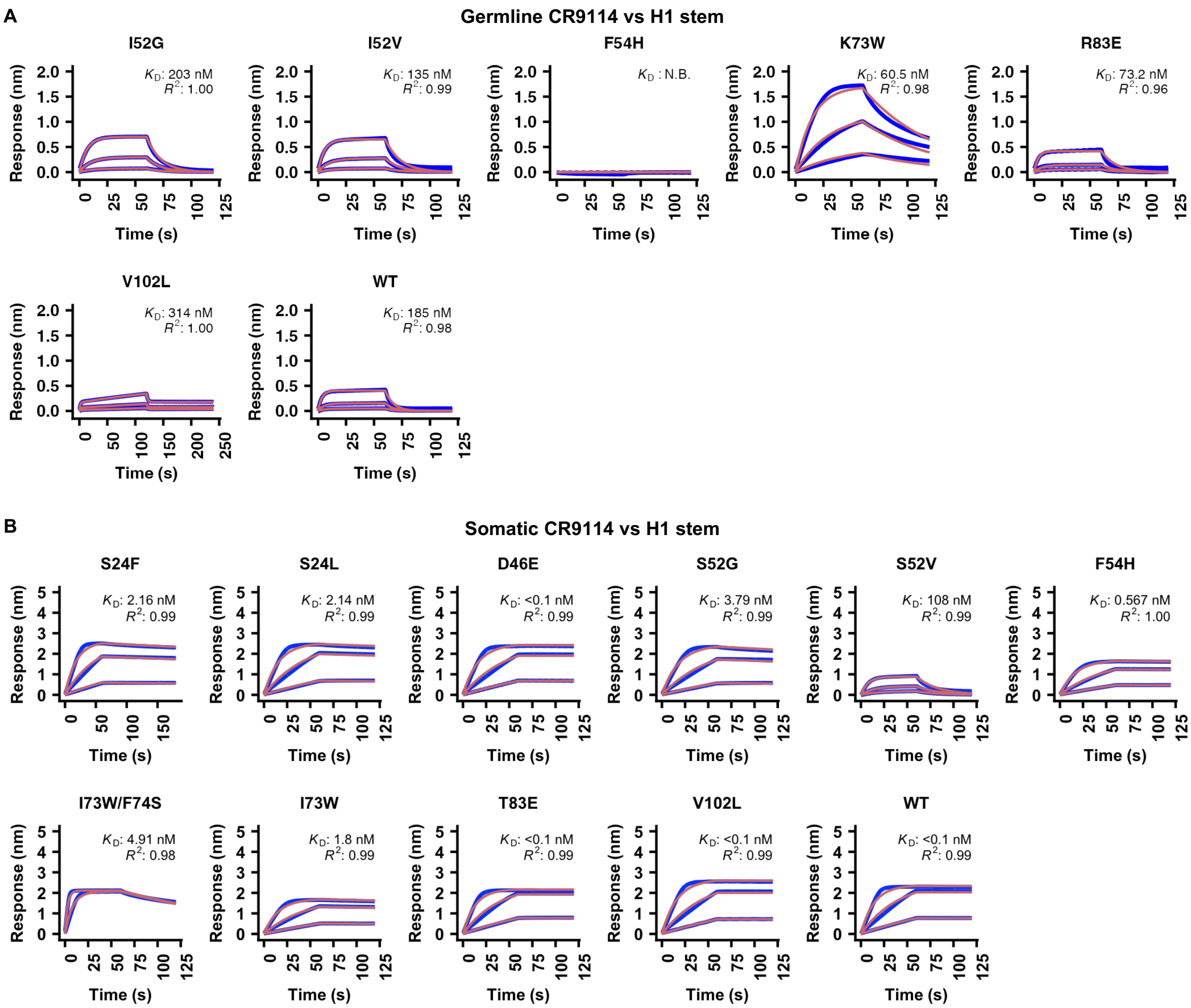


**Figure S3. Binding kinetics of germline and somatic CR9114 mutants against H1 stem.** The binding kinetics of the indicated **(A)** germline and **(B)** somatic CR9114 mutants in Fab format against H1 stem were measured by biolayer interferometry. The Y-axis indicates the signal response. Blue lines represent the response curve, and red lines represent the best fit model (1:1 binding model or 2:1 heterogeneous ligand model, **see Methods**). Binding kinetics were measured for two to three concentrations of Fab at 3-fold dilution. The dissociation constants ($K_D$) and the goodness-of-fit values ($R^2$) are shown.

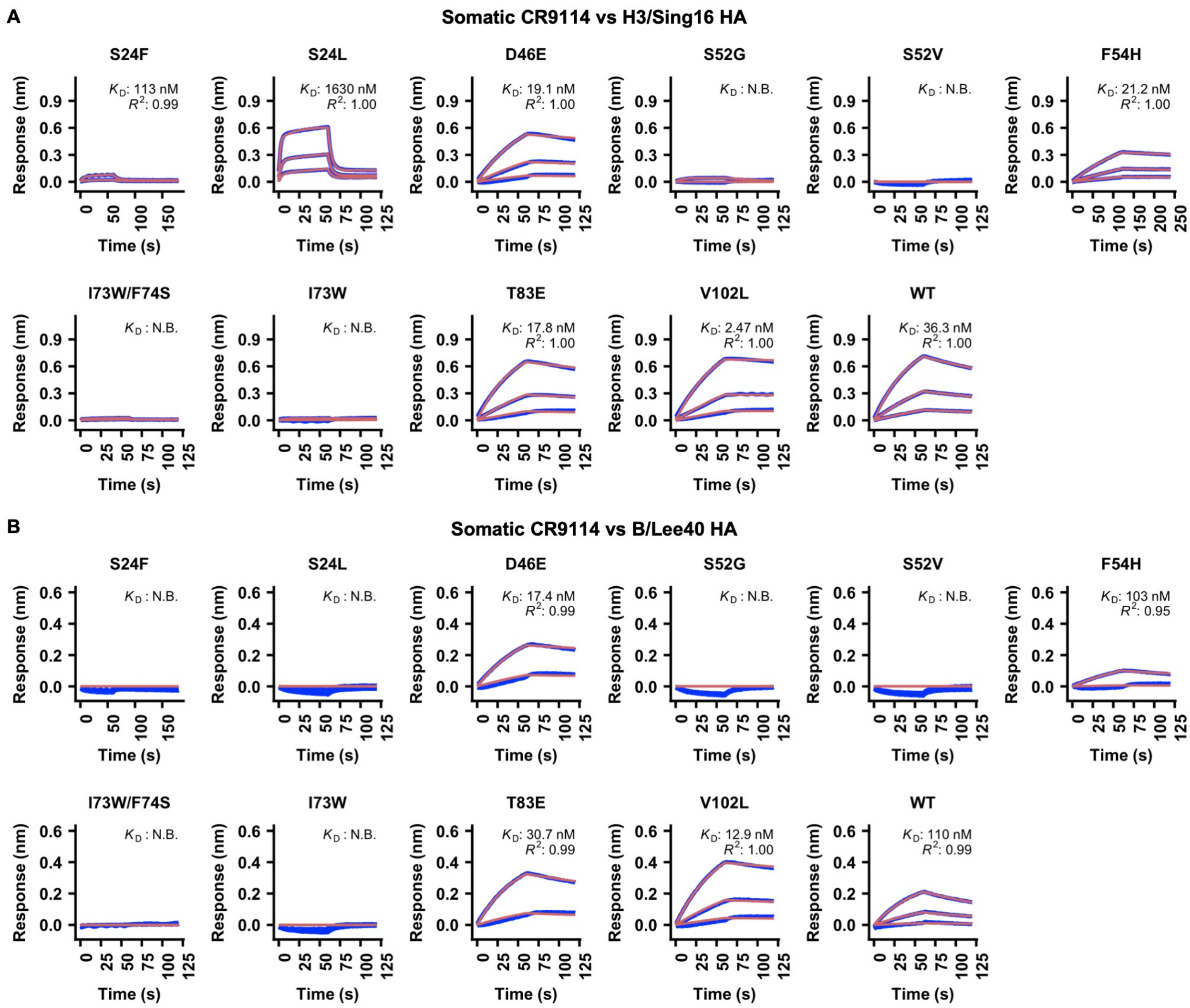


**Figure S4. Binding kinetics of somatic CR9114 mutants against H3 HA and BHA.** The binding kinetics of the indicated somatic CR9114 mutants in Fab format against **(A)** H3 HA and **(B)** BHA were measured by biolayer interferometry. The Y-axis indicates the signal response. Blue lines represent the response curve, and red lines represent the best fit model (1:1 binding model or 2:1 heterogeneous ligand model, **see Methods**). Binding kinetics were measured for two to three concentrations of Fab at 3-fold dilution. The dissociation constants ($K_D$) and the goodness-of-fit values ($R^2$) are shown.

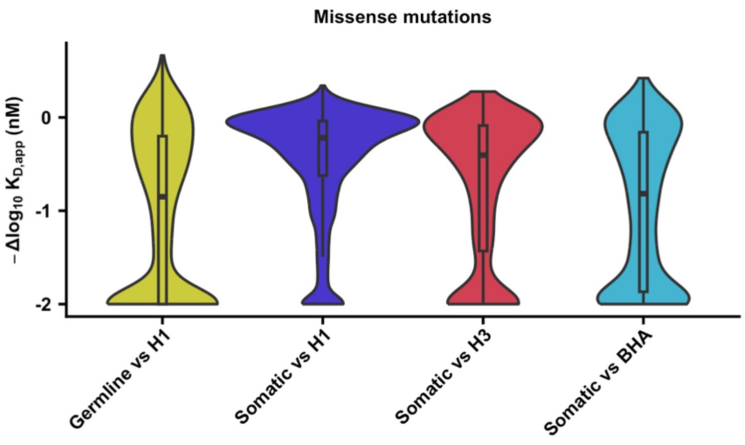


**Figure S5. Distribution of missense mutations effects.** The distribution of -$\Delta\log_{10}$ $K_{D,app}$ values for each deep mutational scanning experiment is shown as a violin plot. “H1”, “H3”, and “BHA” indicate H1 stem, H3/Sing16 HA, and B/Lee40 HA, respectively.

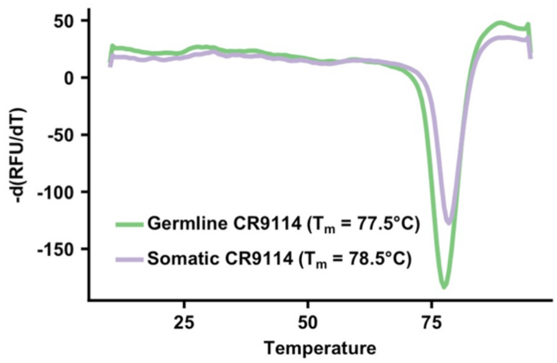


**Figure S6. Measuring the thermal stability of germline and somatic CR9114 through a thermal shift assay.** The first differential curves for the relative fluorescence unit (RFU) with respect to temperature are shown for germline CR9114 (green) and somatic CR9114 (purple). The reported $T_m$ is an average of three independent biological replicates.

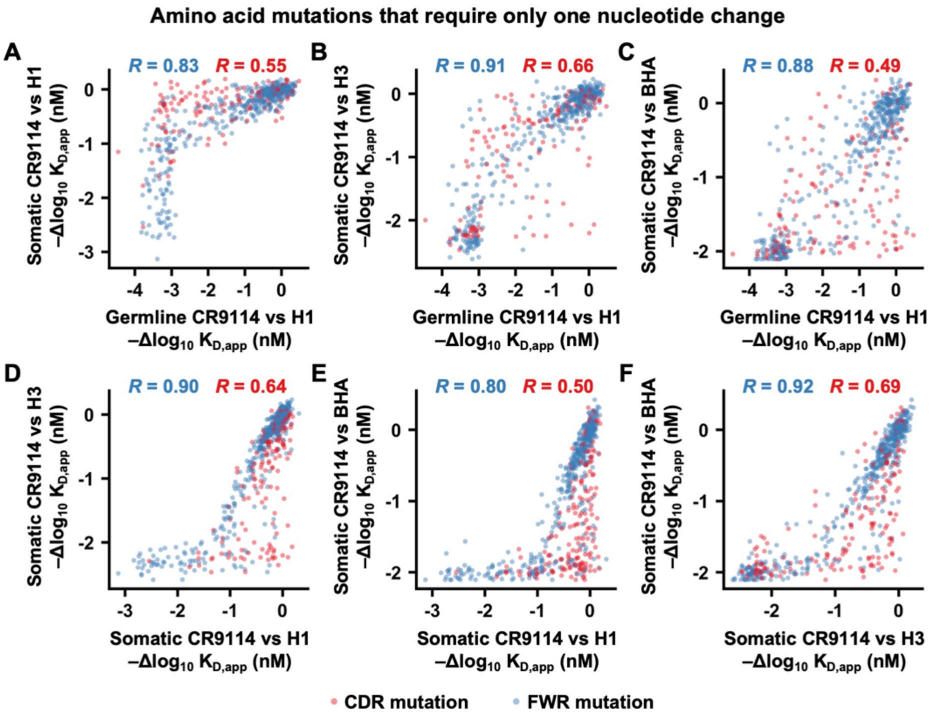


**Figure S7. Correlation of mutational effects on binding affinity across different deep mutational scanning experiments for amino acid mutations that are one nucleotide different from the wild type. (A-F)** The $-\Delta\log_{10} K_{D,app}$ value of each mutation was compared between the deep mutational scanning experiments of **(A)** the germline CR9114 against H1 stem and the somatic CR9114 against H1 stem, **(B)** the germline CR9114 against H1 stem and the somatic CR9114 against H3 HA, **(C)** the germline CR9114 against H1 stem and the somatic CR9114 against BHA, **(D)** the somatic CR9114 against H1 stem and the somatic CR9114 against H3 HA, **(E)** the somatic CR9114 against H1 stem and the somatic CR9114 against BHA, **(F)** the somatic CR9114 against H3 HA and the somatic CR9114 against BHA. Each data point represents a mutation. Pearson correlation coefficients for the mutations in the CDRs and the DE loop (blue) as well as for those in the framework regions (FWRs, red) are indicated. Only those amino acid mutations that are one-nucleotide different from the wild type (i.e. germline or somatic CR9114) are included here. “H1”, “H3”, and “BHA” indicate H1 stem, H3/Sing16 HA, and B/Lee40 HA, respectively.

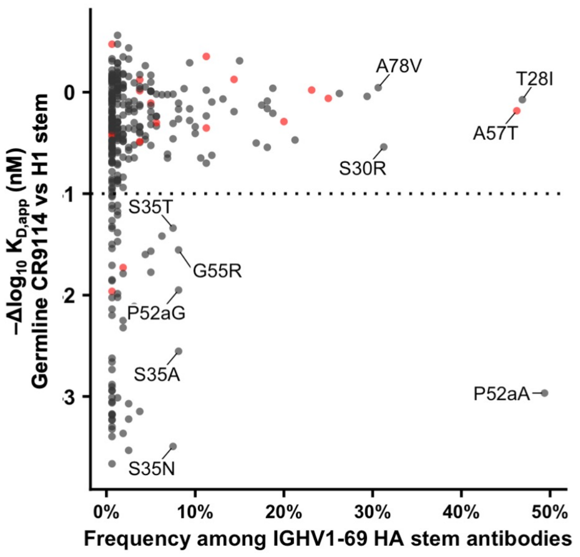


**Figure S8. Correlation between the frequency of somatic hypermutations observed among known IGHV1-69 HA stem antibodies and their effects on germline CR9114 binding to H1 stem.** The occurrence frequency of each somatic hypermutation among the 160 known IGHV1-69 HA stem antibodies[1] is plotted against their corresponding -$\Delta \log_{10} K_{D,app}$ value from the deep mutational scanning experiment of CR9114 against H1 stem. Points in red represent somatic mutations that are present in CR9114.
